# Folate and Tolazamide: Potential competitive inhibitors of barbiturate by in-silico molecular docking

**DOI:** 10.1101/2020.06.27.174938

**Authors:** Kumar Sharp

**Affiliations:** 2^nd^ MBBS undergraduate student, Government Medical College and Hospital, Jalgaon

**Keywords:** barbiturate, toxicity, tolazamide, 5-methyltetrahydrofolate, inhibitor

## Abstract

Barbiturates are the first line drugs for treatment of epilepsy for adults and children in the developing world because of its low cost and proven effectiveness. Due to their adverse effects, no effective management of toxicity and difficulty in determining correct dosage, benzodiazepines are preferred over them. It is also used as a recreational drug, thereby contributing to cases of overdose. My present study was to find the inhibitors of barbiturates by in-silico molecular docking. This approach can highlight only competitive inhibitors as molecules in-silico are rigid and do not produce conformational changes in structure by allosteric binding. 450 FDA-approved drugs were docked to active site of barbiturate of Gleobacter ligand-gated ion channel (GLIC). Drug interactions were visualized, literature search was done to bring out the final results. Tolazamide, an oral anti-diabetic drug and 5-methyltetrahydrofolate, active metabolite of folic acid produced desired results. These results should be used clinically for validation.

## Introduction

Barbiturates are the first line drugs for treatment of epilepsy for adults and children in the developing world because of its low cost and proven effectiveness. Due to their adverse effects, no effective management of toxicity and difficulty in determining correct dosage, benzodiazepines are preferred over them. It is also used as a recreational drug, thereby contributing to cases of overdose ^[1]^. My present study was to find the inhibitors of barbiturates by in-silico molecular docking. This approach can highlight only competitive inhibitors as molecules in-silico are rigid and do not produce conformational changes in structure by allosteric binding ^[2]^.

## Methodology

Gleobacter ligand-gated ion channel (GLIC) tertiary structure 5L47 ^[3]^ was downloaded from Protein Data Bank (Figure 1). FDA-approved drug structures were downloaded from Zinc15 database ^[4]^ consisting of around 450 compounds. Selenocyanobarbital was the desired ligand in GLIC structure (Figure 2). The target active site to which selenocyanobarbital binds was visualized using Drug Discover Studio ^[5]^. After determining the amino acids of the target active site, the structure was cleaned of water molecules, ligands and other heterogenous atoms or molecules. Thiopental was chosen as the reference barbiturate molecule. Its three-dimensional structure was downloaded from Zinc15 database. PyRx software ^[6]^ was used for molecular docking because of its graphical user interface and inclusion of two important software, Autodock Vina ^[7]^ and Open Babel ^[8]^. The operations were performed on a Windows 10 operating system utilizing 8 processors. All drug structures obtained from Zinc15 were processed in Open Babel to convert them to the minimum energy conformation. Partial charges and hydrogen were added and the structures were converted into Autodock ligands. The GLIC structure was then processed into Autodock molecule using PyRx. Autodock Vina wizard was used to dock thiopental to the active site determined before. The search space was adjusted in three-dimension to avoid undesirable binding. Exhaustiveness was kept at the maximum of 8. Of the 8 modes in which thiopental bonded, the first mode which had root mean square deviation (RMSD) of 0.00 was chosen as reference. The binding affinity of this mode E_0_ was chosen as the reference binding energy. In the same way, all the FDA-approved drug structures underwent molecular docking. Their binding affinity with RMSD equal to 0.00 was compiled. Binding affinity (kcal/mol) in negative signifies stronger binding. Hence, more the value in negative, more is the binding affinity. Therefore top 35 ligands which had a more binding affinity than thiopental E_0_ are chosen for deep analysis. The ligands/drugs which are banned for use in India were removed from the list. Since ligands can interact with other amino acids in search space other than the target amino acids, these interactions with the receptor/GLIC was visualized using Discovery Studio. The number of desired interactions achieved were noted and then the results were sorted to provide the final candidates which interacted with at least 4 of the 6 amino acids. All the above steps taken improve the accuracy of the results which is desired.

**Figure 1:**
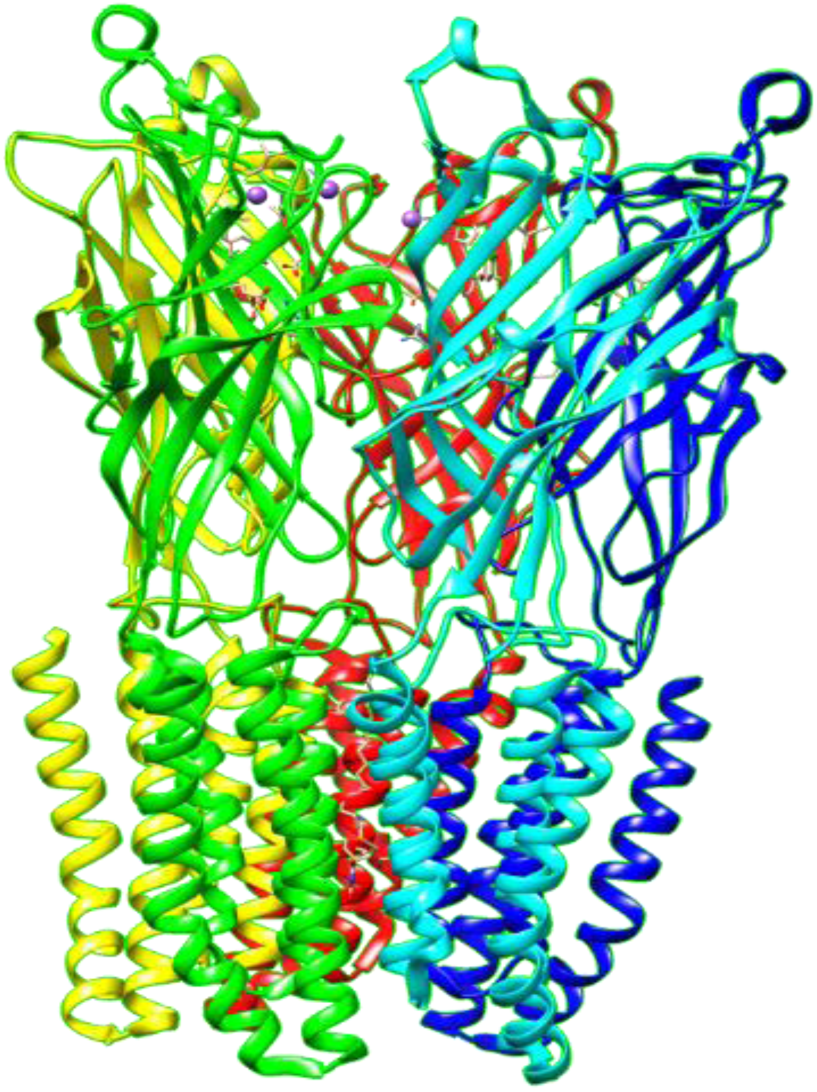
Gleobacter ligand-gated ion channel (GLIC): 5L47 from Protein Data Bank. Different colours represent different chains.

**Figure 2:**
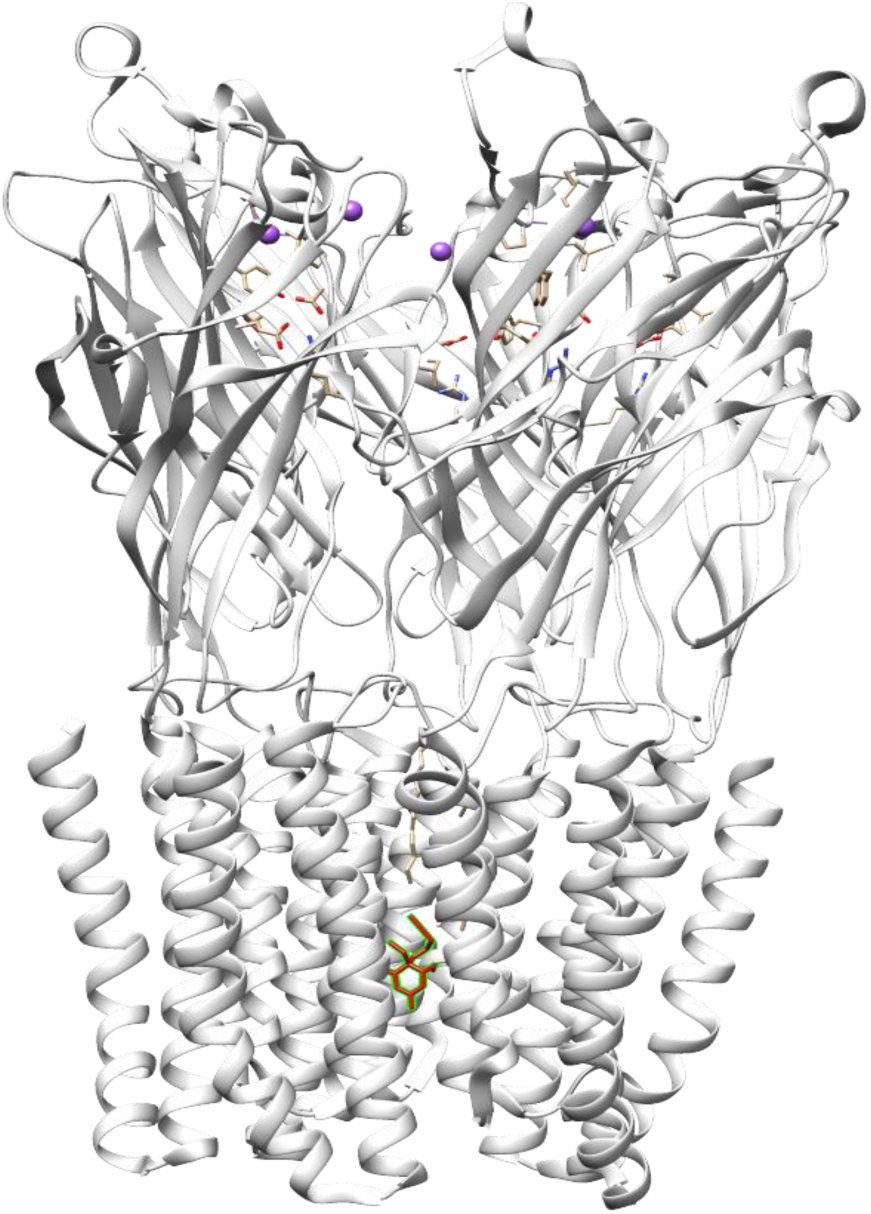
Selenocyanobarbital (red) bound to active site in GLIC (PDB ID:5L47)

## Results

There were 6 active site amino acids determined from the interaction between Selenocyanobarbital and GLIC. These are listed as follows and represented in the diagram (Figure 3):

**Figure 3:**
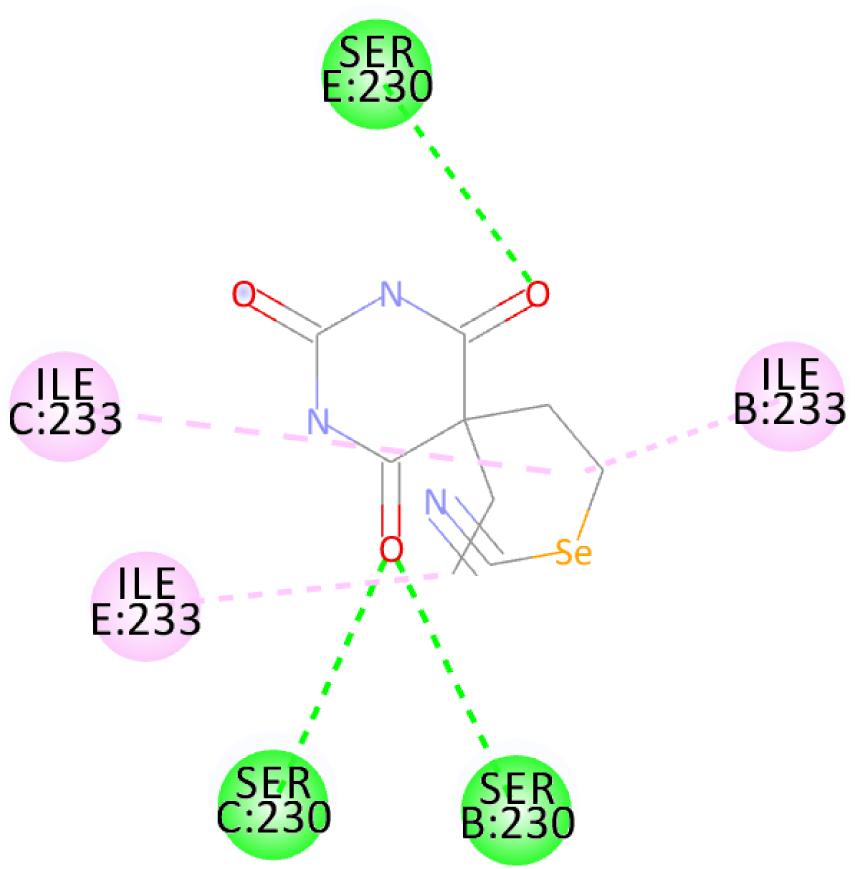
Selenocyanobarbital interactions with GLIC amino acids. Conventional hydrogen bonds are marked in green and alkyl bonds are marked in pink.

1. SER chain B:230
2. ILE chain B:233
3. SER chain C:230
4. ILE chain C:233
5. SER chain E:230
6. ILE chain E:233

ILE and SER are standard abbreviations for amino acids isoleucine and serine respectively. Numbers indicate their position in the chain. Active sites in GLIC are selected for docking in PyRx. Autodock Vina search space was restricted to the following co-ordinates taking care to included all active sites. The search space co-ordinates were as follows:

center_x = 60.2389306218

center_y = -26.5411159502

center_z = 54.8963565848

size_x = 15.4114621461

size_y = 13.6281669465

size_z = 13.8838945818

E_0_ binding affinity of thiopental was −6.1 kcal/mol. On the basis of the conditions set above, the final set of ligands obtained was as follows (Table 1):

**Table 1:**
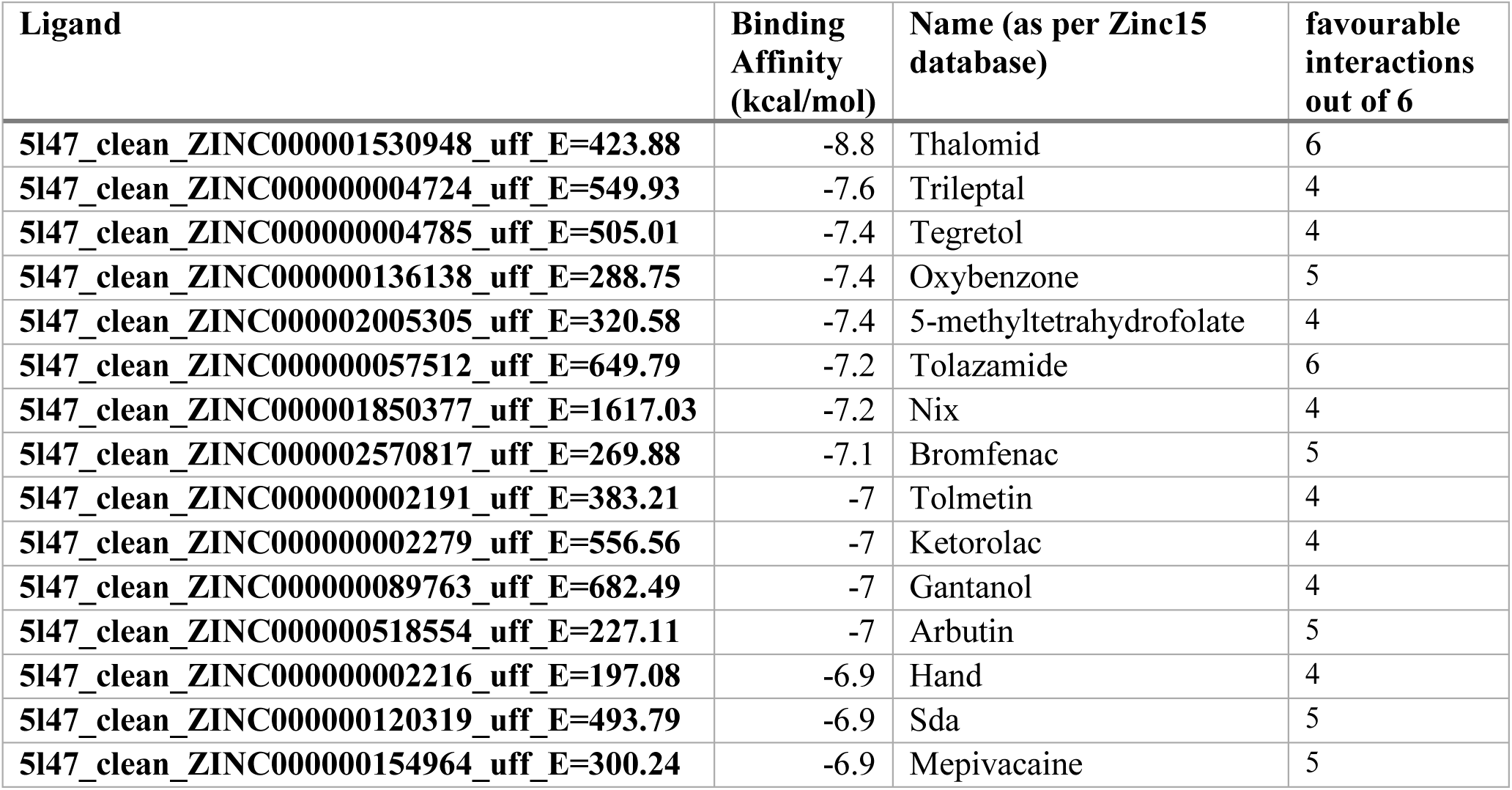
Top 15 ligands arranged in increasing order of binding affinity along with the number of desired interactions greater than or equal to 4.

These interactions are represented in the 2-dimensional diagram below (Figure 4-18)

**Figure 4:**
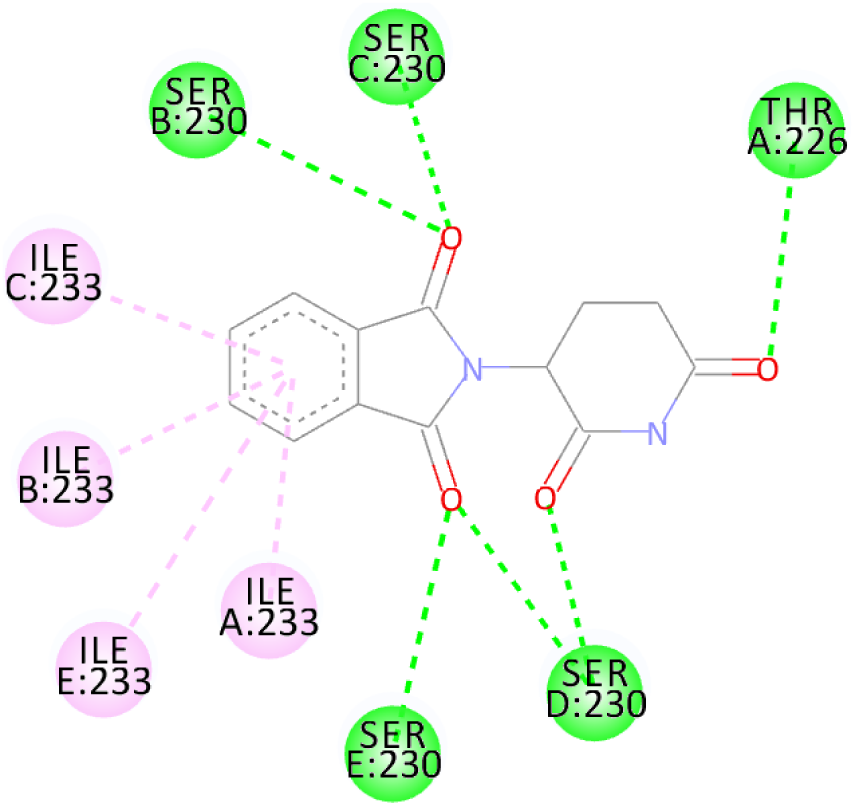
Thalomid interaction with GLIC receptor amino acids (shown in circles). Conventional hydrogen bonds are represented in green and pi-alkyl bonds are represented in pink.

**Figure 5:**
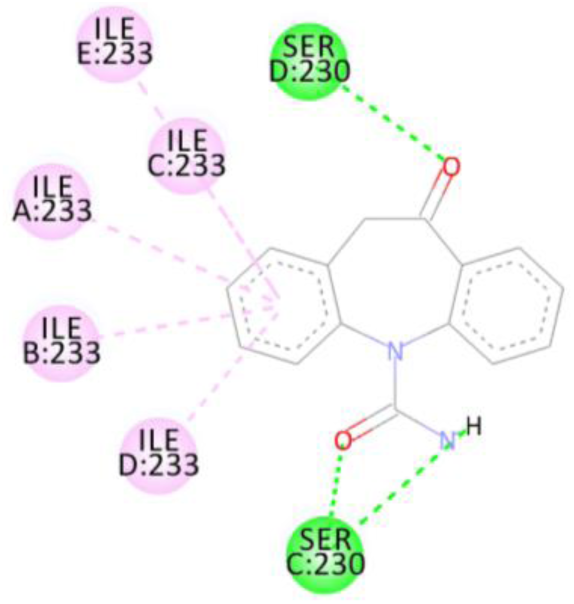
Trileptal interaction with GLIC receptor amino acids (shown in circles). Conventional hydrogen bonds are represented in green and pi-alkyl bonds are represented in pink.

**Figure 6:**
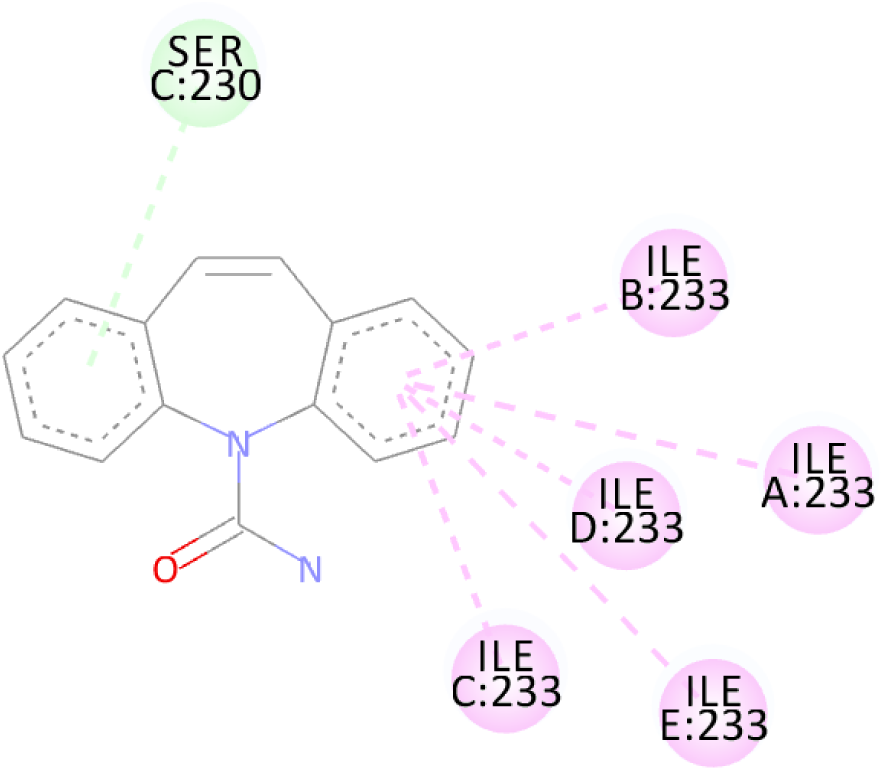
Tegretol interaction with GLIC receptor amino acids (shown in circles). Pi-donor hydrogen bonds are represented in light blue/ light green and pi-alkyl bonds are represented in pink.

**Figure 7:**
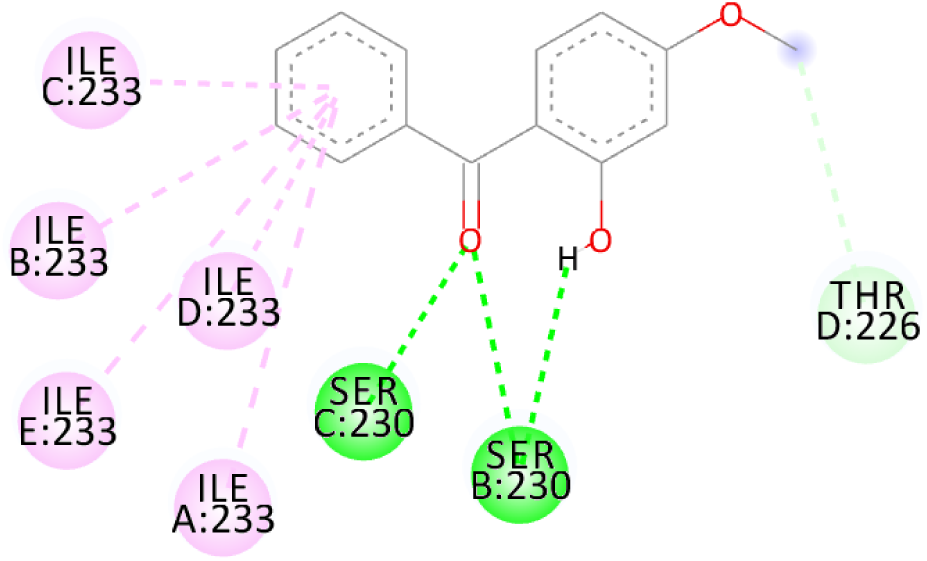
Oxybenzone interaction with GLIC receptor amino acids (shown in circles). Carbon hydrogen bonds are represented in light blue/ light green, pi-alkyl bonds are represented in pink and conventional hydrogen bonds in green.

**Figure 8:**
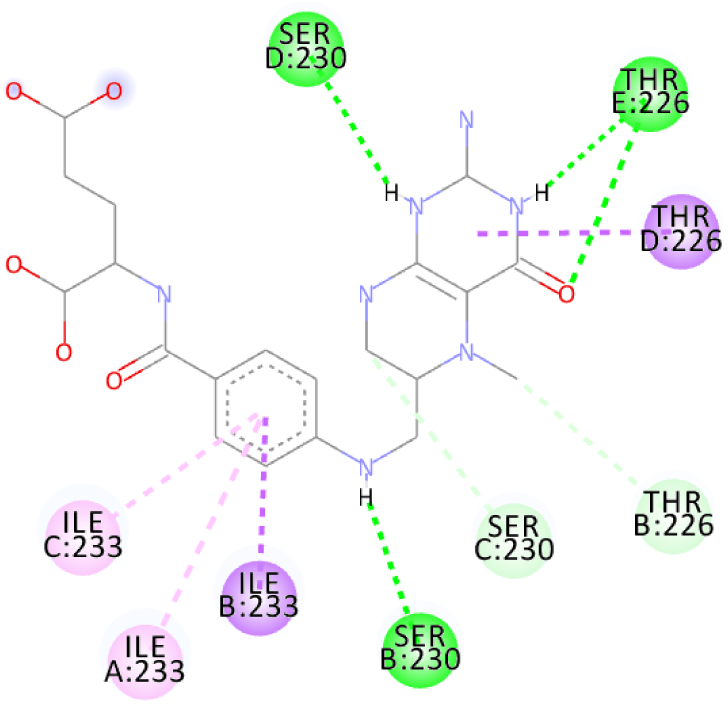
5-methyltetrahydrofolate interaction with GLIC receptor amino acids (shown in circles). Carbon hydrogen bonds are represented in light blue/ light green, pi-sigma bonds in purple, pi-alkyl bonds are represented in pink and conventional hydrogen bonds in green.

**Figure 9:**
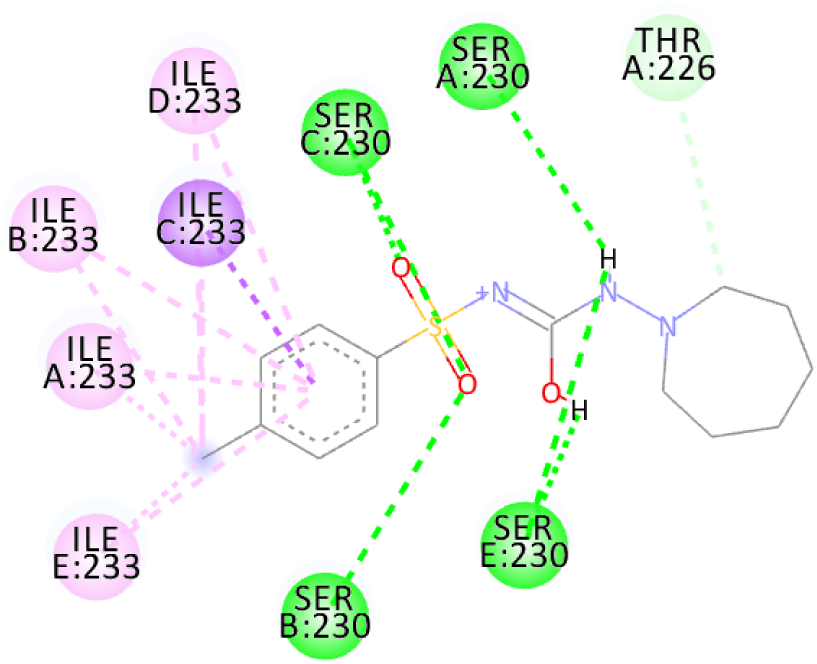
Tolazamide interaction with GLIC receptor amino acids (shown in circles). Carbon hydrogen bonds are represented in light blue/ light green, pi-sigma bonds in purple, pi-alkyl bonds are represented in pink and conventional hydrogen bonds in green.

**Figure 10:**
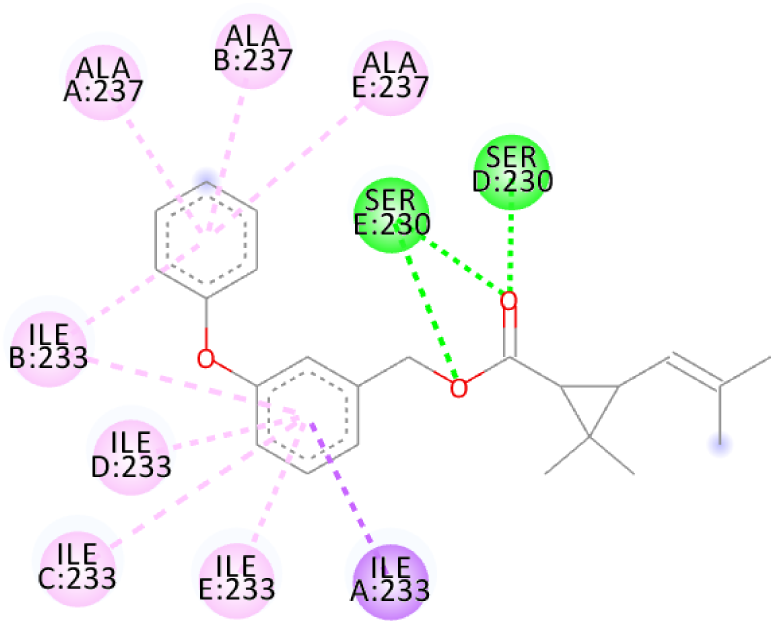
Nix interaction with GLIC receptor amino acids (shown in circles). Pi-sigma bonds are represented in purple, pi-alkyl bonds are represented in pink and conventional hydrogen bonds in green.

**Figure 11:**
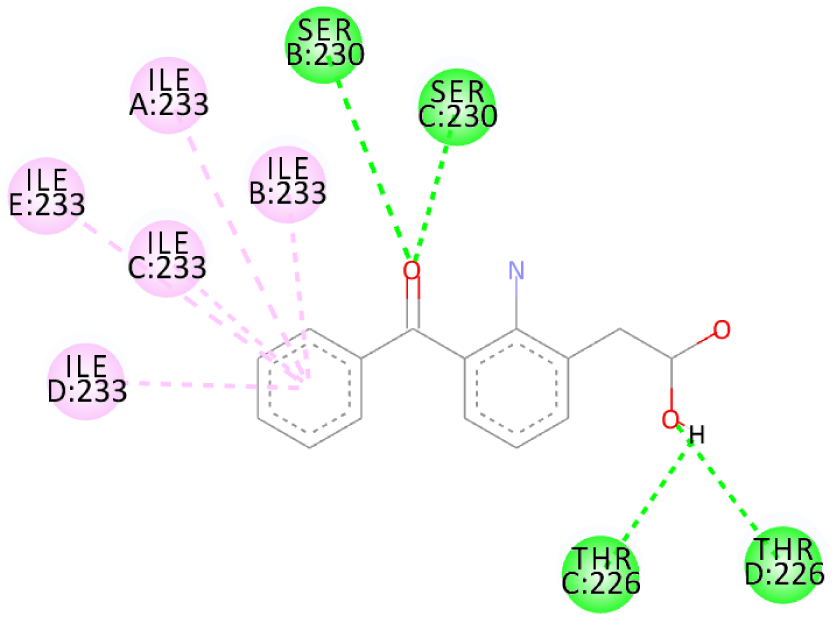
Bromfenac interaction with GLIC receptor amino acids (shown in circles). Conventional hydrogen bonds are represented in green and pi-alkyl bonds are represented in pink.

**Figure 12:**
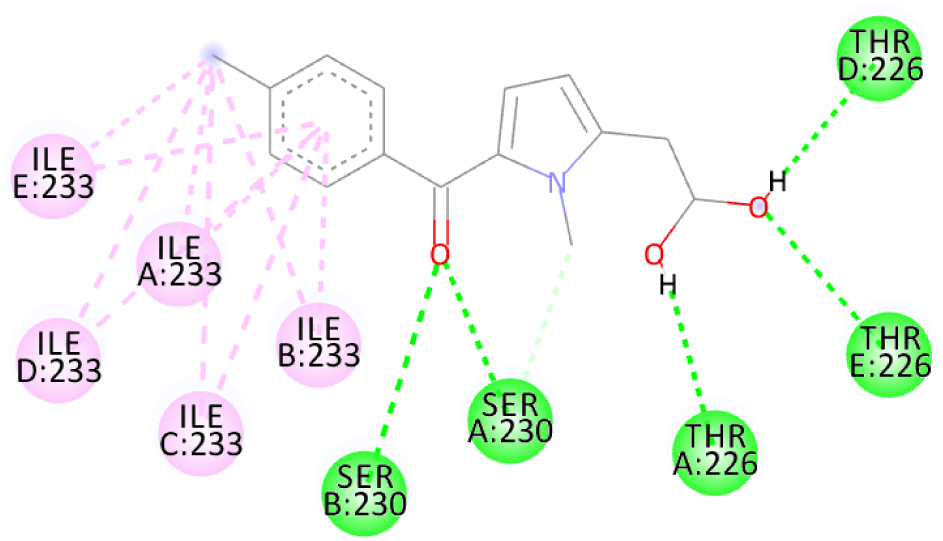
Tolmetin interaction with GLIC receptor amino acids (shown in circles). Conventional hydrogen bonds are represented in green and pi-alkyl bonds are represented in pink.

**Figure 13:**
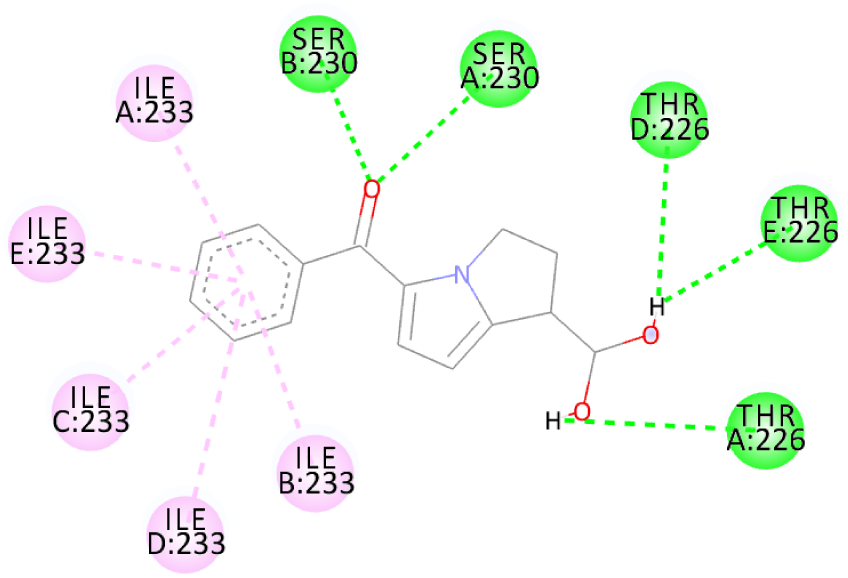
Ketorolac interaction with GLIC receptor amino acids (shown in circles). Conventional hydrogen bonds are represented in green and pi-alkyl bonds are represented in pink.

**Figure 14:**
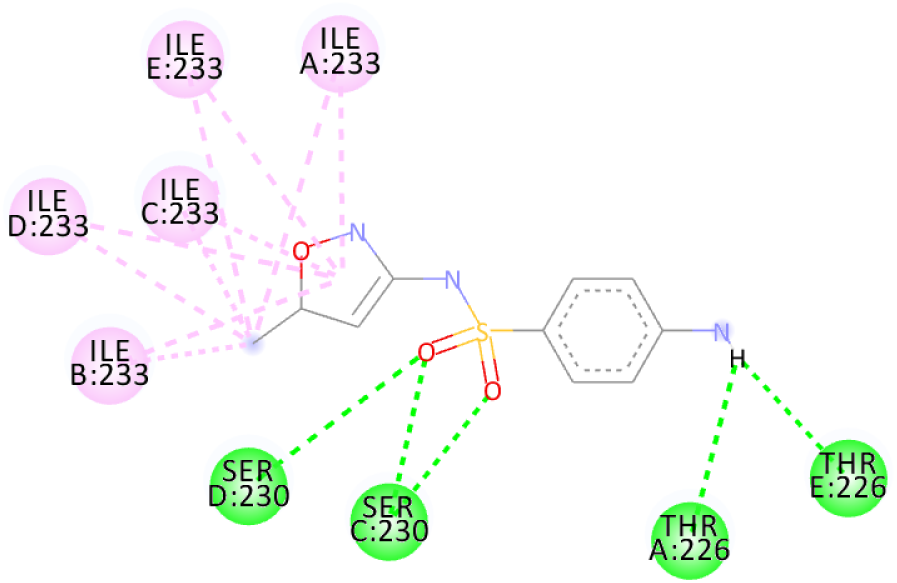
Gantanol interaction with GLIC receptor amino acids (shown in circles). Conventional hydrogen bonds are represented in green and pi-alkyl bonds are represented in pink.

**Figure 15:**
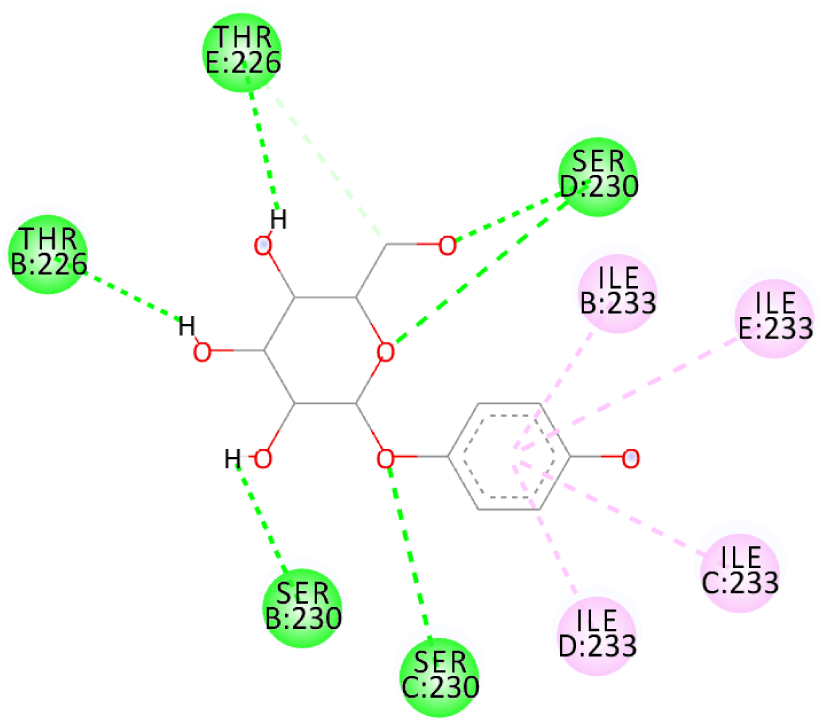
Arbutin interaction with GLIC receptor amino acids (shown in circles). Conventional hydrogen bonds are represented in green and pi-alkyl bonds are represented in pink.

**Figure 16:**
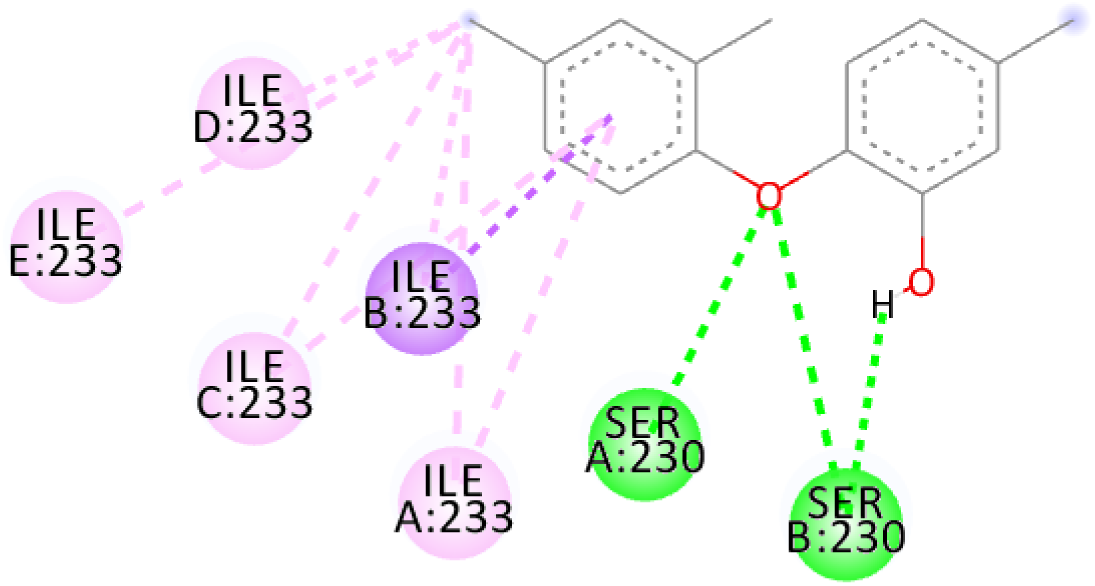
Hand interaction with GLIC receptor amino acids (shown in circles). Pi-sigma bonds are represented in purple, pi-alkyl bonds are represented in pink and conventional hydrogen bonds in green.

**Figure 17:**
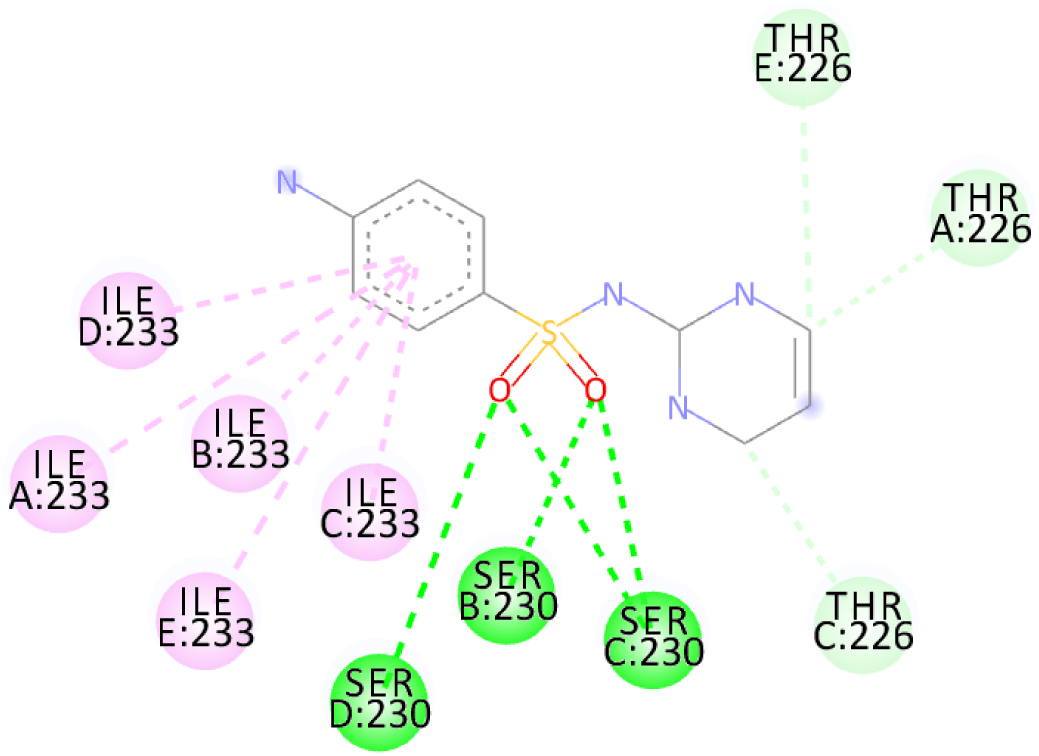
Sda interaction with GLIC receptor amino acids (shown in circles). Carbon hydrogen bonds are represented in light blue/ light green, pi-alkyl bonds are represented in pink and conventional hydrogen bonds in green.

**Figure 18:**
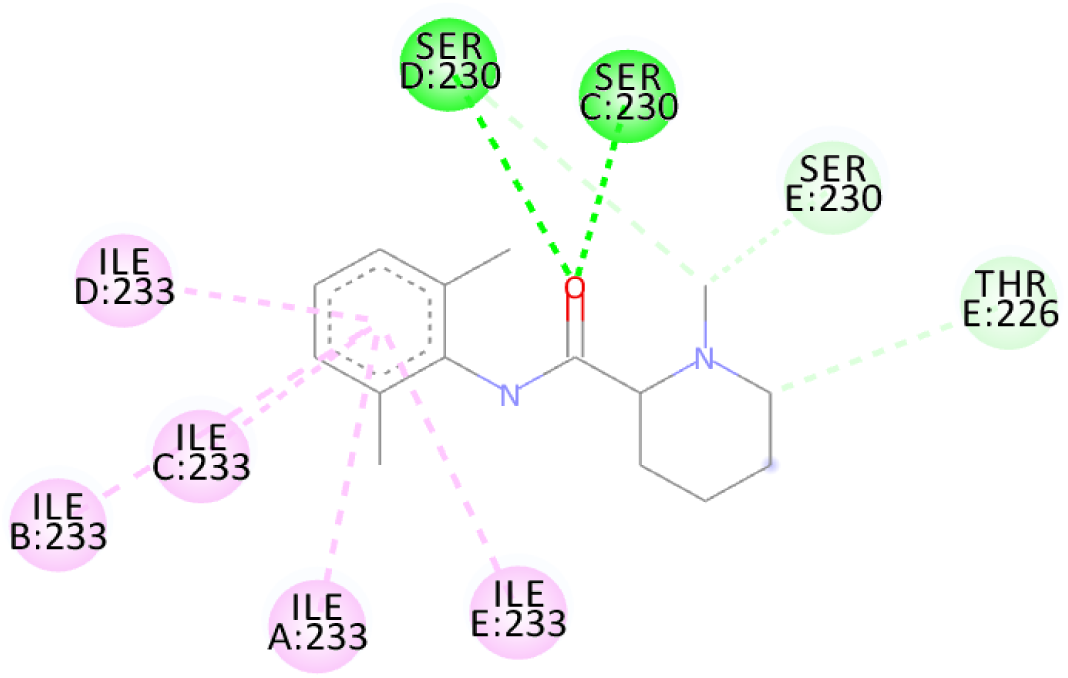
Mepivacaine interaction with GLIC receptor amino acids (shown in circles). Carbon hydrogen bonds are represented in light blue/ light green, pi-alkyl bonds are represented in pink and conventional hydrogen bonds in green.

## Discussion

The main objective of this study is to find an agent which can reverse the effect the barbiturate. Literature search for all the above compounds were done to find any documented relation between them and barbiturates. Thalomid which is the trade name for Thalidomide is used as a non-barbiturate sedative. It also enhances the effects of barbiturate ^[9]^ and hence is against the objective. Trileptal and tegretol are trade names for oxcarbazepine and carbamazepine respectively. They are used as seizure control drugs ^[10][11]^ and preferred over barbiturate but in no way can reverse the effect of barbiturate as they have the same clinical action. Tolazamide which is an oral anti-diabetic drug has been documented to interfere with barbiturate ^[12][13]^. This is highly probable due to the fact that they both target similar active site found in this study. It can hence serve as competitive inhibitor of barbiturates. 5-methyltetrahydrofolate which is the active metabolite of folic acid has been experimented upon in the past and has shown inverse relation with phenytoin and barbiturate level in the body ^[14]^. Patients receiving anti-convulsant therapy consisting of phenytoin and barbiturate had developed megaloblastic anaemia due to folate deficiency ^[15]^. Gantanol which is trade name for sulfamethoxazole had only one study with relation to barbiturates ^[16]^. However, that study could not be accessed. No relevant literature could be found for other compounds of this study. The final two candidates after this study are tolazamide and 5-methyltetrahydrofolate. These drugs can be used in future for clinical trials for management of barbiturate overdose.

## Conclusion

The above study has brought out probable treatment of barbiturate overdose as tolazamide and 5-methyltetrahydrofolate. Antidote for drugs need to exist as long as they are used for public. Clinical trials are however necessary to validate these results. Non-competitive inhibitors have to be found by in-vitro trials as it is the most important limitation of in-silico approach.

## Ethical consideration

Not required

## References

1. Katzung BG. Basic and clinical pharmacology. Mc Graw Hill; 2012.

2. Sharp K, Dange DS. In-Silico FDA-Approved Drug Repurposing to Find the Possible Treatment of Coronavirus Disease-19 (COVID-19) [Internet]. ChemRxiv; 2020 [cited 2020Jun27]. Available from: https://chemrxiv.org/articles/In-Silico_FDA-Approved_Drug_Repurposing_to_Find_the_Possible_Treatment_of_Coronavirus_Disease-19_COVID-19_/12340718/1

3. Fourati Z, Ruza RR, Laverty D, Drège E, Delarue-Cochin S, Joseph D, Koehl P, Smart T, Delarue M. Barbiturates bind in the GLIC ion channel pore and cause inhibition by stabilizing a closed state. Journal of Biological Chemistry. 2017 Feb 3;292(5):1550–8.

4. Sterling T, Irwin JJ. ZINC 15–ligand discovery for everyone. Journal of chemical information and modeling. 2015 Nov 23;55(11):2324–37.

5. Ref. Dassault Systèmes BIOVIA, Discovery Studio Modeling Environment, Release 2017, San Diego: Dassault Systèmes, 2016.

6. Dallakyan S, Olson AJ. Small-molecule library screening by docking with PyRx. InChemical biology 2015 (pp. 243–250). Humana Press, New York, NY.

7. Trott O, Olson AJ. AutoDock Vina: improving the speed and accuracy of docking with a new scoring function, efficient optimization, and multithreading. Journal of computational chemistry. 2010 Jan 30;31(2):455–61.

8. O’Boyle NM, Banck M, James CA, Morley C, Vandermeersch T, Hutchison GR. Open Babel: An open chemical toolbox. Journal of cheminformatics. 2011 Dec;3(1):33.

9. Laffitte E, Revuz J. Thalidomide: an old drug with new clinical applications. Expert opinion on drug safety. 2004 Jan 1;3(1):47–56.

10. Schmidt D, Arroyo S, Baulac M, Dam M, Dulac O, Friis ML, Kälviäinen R, Krämer G, Van Parys J, Pedersen B, Sachdeo R. Recommendations on the clinical use of oxcarbazepine in the treatment of epilepsy: a consensus view. Acta neurologica scandinavica. 2001 Sep;104(3):167–70.

11. Shorvon SD, Reynolds EH. Anticonvulsant peripheral neuropathy: a clinical and electrophysiological study of patients on single drug treatment with phenytoin, carbamazepine or barbiturates. Journal of Neurology, Neurosurgery & Psychiatry. 1982 Jul 1;45(7):620–6.

12. Thürkow I, Wesseling H, Meijer DK. Estimation of phenytoin in body fluids in the presence of sulphonyl urea compounds. Clinica Chimica Acta. 1972 Mar 1; 37:509–13.

13. Logie AW, Galloway DB, Petrie JC. Drug interactions and long-term antidiabetic therapy. British journal of clinical pharmacology. 1976 Dec;3(6):1027–32.

14. Elsborg L. Binding of folic acid to human plasma proteins. Acta Haematologica. 1972;48(4):207–12.

15. Klipstein FA. Subnormal serum folate and macrocytosis associated with anticonvulsant drug therapy. Blood. 1964 Jan;23(1):68–86.

16. Kerr JC, Lazaro EJ, Cohen IS, Spillert CR, Devereux C. Potentiating effect of trimethoprim and sulfamethoxazole on barbiturate anesthesia. InCLINICAL RESEARCH 1984 Jan 1 (Vol. 32, No. 3, pp. A706-A706). 6900 GROVE RD, THOROFARE, NJ 08086: SLACK INC.

